# Ecological inference using data from accelerometers needs careful protocols

**DOI:** 10.1101/2021.07.07.451487

**Authors:** Baptiste Garde, Rory P. Wilson, Adam Fell, Nik Cole, Vikash Tatayah, Mark D. Holton, Kayleigh A. R. Rose, Richard S. Metcalfe, Hermina Robotka, Martin Wikelski, Fred Tremblay, Shannon Whelan, Kyle H. Elliott, Emily L. C. Shepard

## Abstract

1. Accelerometers in animal-attached tags have proven to be powerful tools in behavioural ecology, being used to determine behaviour and provide proxies for movement-based energy expenditure. Researchers are collecting and archiving data across systems, seasons and device types. However, in order to use data repositories to draw ecological inference, we need to establish the error introduced according to sensor type and position on the study animal and establish protocols for error assessment and minimization.
2. Using laboratory trials, we examine the absolute accuracy of tri-axial accelerometers and determine how inaccuracies impact measurements of dynamic body acceleration (DBA), as the main acceleration-based proxy for energy expenditure. We then examine how tag type and placement affect the acceleration signal in birds using (i) pigeons *Columba livia* flying in a wind tunnel, with tags mounted simultaneously in two positions, (ii) back- and tail-mounted tags deployed on wild kittiwakes *Rissa tridactyla*. Finally, we (iii) present a case study where two generations of tag were deployed using different attachment procedures on red-tailed tropicbirds *Phaethon rubricauda* foraging in different seasons.
3. Bench tests showed that individual acceleration axes required a two-level correction (representing up to 4.3% of the total value) to eliminate measurement error. This resulted in DBA differences of up to 5% between calibrated and uncalibrated tags for humans walking at different speeds. Device position was associated with greater variation in DBA, with upper- and lower back-mounted tags in pigeons varying by 9%, and tail- and back-mounted tags varying by 13% in kittiwakes. Finally, DBA varied by 25% in tropicbirds between seasons, which may be attributable to tag attachment procedures.
4. Accelerometer accuracy, tag placement, and attachment details critically affect the signal amplitude and thereby the ability of the system to detect biologically meaningful phenomena. We propose a simple method to calibrate accelerometers that should be used prior to deployments and archived with resulting data, suggest a way that researchers can assess accuracy in previously collected data, and caution that variable tag placement and attachment can increase sensor noise and even generate trends that have no biological meaning.

## Introduction

Animal-attached tags have revolutionized our understanding of wild animal ecology (Bograd et al., 2010; Sequeira et al., 2021; Yoda, 2019). Of the sensors often used, accelerometers (Yoda et al., 1999) are regarded as particularly powerful tool for studying wild animal behavioural ecology, with studies using them to look at the occurrence and intensity of behaviour (Chakravarty et al., 2019; Fehlmann et al., 2017), assess movement characteristics (Shepard et al., 2008) and as a proxy for energy expenditure (Wilson et al., 2020). The latter has developed rapidly since the demonstration that dynamic body acceleration (DBA) is related to energy expenditure across a range of vertebrates and invertebrates (Halsey et al., 2009; Wilson et al., 2019, 2006). Such measurements have great potential for understanding animal strategies, in particular studying how animals respond to change in food availability (Kokubun et al., 2011), climate (Gudk et al., 2019) and anthropogenic threats or activity (Nickel et al., 2021; Payne et al., 2015; Yorzinski et al., 2015).

In mammals, accelerometers tend to be attached using collars, and while collars have their own complications in terms of the need to obtain a good fit and account for collar rotation in data interpretation (Wilson et al., 2020), the position of attachment is largely standardised. In contrast, researchers use different attachment positions on birds. For instance, tags are deployed on the lower back, the tail or the belly of seabirds depending on the species and the tag position associated with least detriment (Elliott, 2016; Ropert-Coudert et al., 2003; Vandenabeele et al., 2014). While some species appear to show less of a response to tags mounted on the back, there are lower weight limits for what can be attached to the tail, and both positions impact flight forces. Researchers working with raptors may deploy tags using backpack or leg-loop harnesses (e.g. Harel et al., 2017; Williams et al., 2015, respectively), which results in differences in tag position. The widespread availability and use of accelerometers means that large datasets, collected over years, are now available, providing valuable information about behaviour including flight effort across temporal and spatial scales (Kranstauber et al., 2011). Unsurprisingly, these data have been collected using different methods of attachment and by deploying a variety of different tags without critical analysis of the compatibility of different datasets (Sequeira et al., 2021).

Tag position on the body is likely to affect acceleration values, as pointed out by Wilson et al. (2020), who noted that DBA (Qasem et al., 2012) varied with tag position in humans wearing back and waist-mounted tags running on a treadmill (with DBA values varying by ~ 0.25 g at intermediate speeds). This is easy to understand since humans have a flexible spine. Birds, on the other hand, have an essentially immoveable box-like thorax (Baumel, 1993). Differences in acceleration between tags placed on the back and the neck (Kölzsch et al., 2016) or the tail (Elliott, 2016) are easy to associate with independent movement of the head or tail, but the thorax itself can experience pitch changes over the wing beat cycles (Su et al., 2012; Tobalske & Dial, 1996), which may affect the acceleration recorded by loggers depending on their position. As part of that, we note that the precise position of the accelerometer chips on the circuit boards may also affect the acceleration measured by the sensors, particularly in cases where the circuit board is long relatively to the bird’s back and where the chip could be positioned close to either end.

At a more fundamental level, the fabrication of loggers with accelerometers involves extensive heating as the sensors are soldered to the circuit boards. This is known to change their sensory performance (output *versus* acceleration) (Ruzza et al., 2018), even if they are carefully calibrated prior to this process (see https://www.mouser.fr/datasheet/2/389/lsm303dlhc-955106.pdf). Specifically, while the vector sum of the 3 acceleration channels should be 1 when a unit is at rest, this can vary after heating, resulting in error in the estimation of the Earth’s gravitational component. This can in turn introduce error into the estimation of the “dynamic” acceleration, or acceleration due to movement, which is the basis for acceleration-based proxies for the energy expenditure (Wilson et al., 2020).

In this manuscript, we assess the error associated with the sensors themselves and how the position and fixing of the accelerometer on the study animal affects acceleration metrics before proposing solutions to minimize these issues. Specifically, we first examine how variability in VeDBA relates to improperly calibrated tri-axial accelerometers, using a case with humans walking defined courses at fixed speeds. We then examine how tag position affects VeDBA and signal amplitude using pigeons (*Columba livia*) flying in a wind tunnel with two tags placed on different locations of their back. Finally, we examine two examples of variation in the acceleration signal based on retrospective analysis of field studies involving; (1) red-tailed tropicbirds (*Phaethon rubricauda*) equipped with two different types of loggers attached using marginally different protocols in two separate seasons, and (2) black-legged kittiwakes (*Rissa tridactyla*) equipped with a tag on the back and one on the tail, as two positions favoured by seabird researchers for tag placement.

## Methods

### Measurement of acceleration accuracy of tri-axial sensors

We first calibrated tri-axial accelerometers within 5 Daily Diary tags (inch board) (Wildbyte Technologies, Swansea University, UK) (Wilson et al., 2008), by setting them motionless on a table in a series of defined orientations (each for ca. 10 seconds). Six orientations (hereafter the ‘6-O method’) were chosen so that the tags always had one of their three acceleration axes perpendicular to gravity and these were rotated according to the 6 axes of a die so that each of the 3 accelerometer axes nominally reads −1 *g* and 1 *g*.

The outputs of these motionless calibrations were then used to derive the six respective maxima of the acceleration vectorial sum given by;

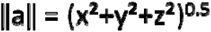

where x, y and z are the raw acceleration values, for the periods when they were held still. Note that there are 6 maxima because each axis has two values: a minimum and a maximum, which become positive in the vectorial sum. In a device with perfect acceleration sensors, all maxima should be 1.0 *g* (although the acceleration on earth varies with latitude by up to a maximum of 0.0053 *g* due to the earth’s shape and the centrifugal force generated by the planet spinning as well as other processes (Novák, 2010). However, values were always either marginally higher or lower than 1.0 *g* (see Results). Furthermore, the two maxima for each axis differed. This therefore requires 2 steps to be corrected, where (1) a correction factor is applied to the lower value to ensure both “maxima” are the same and then (2) the same offset is applied to both readings to convert readings to exactly 1.0 *g*.

Subsequently, tags were deployed on 12 people, attached to the lower back using elastic. Each person walked back and forth on a 25 m straight-line course at four different speeds (0.69, 0.97, 1.25 and 1.53 m s^−1^; randomly ordered), each for 3 minutes. Speeds were held constant using a metronome. The mean VeDBA (defined as VeDBA = (x_D_^2^+y_D_^2^+z_D_^2^)^0.5^ where x_D_, y_D_ and z_D_ are the dynamic body acceleration recorded by each of the three channels of acceleration – for details see Wilson et al. (2020), was calculated across each 3-minute trial with, and without, the calibration corrections.

### Effect of tag position on acceleration

The effect of tag position was first tested on three pigeons (*Columba livia*) flying under controlled conditions in a wind tunnel at speeds ranging from 10 to 22 m.s^−1^. Birds were equipped simultaneously with two tags recording acceleration at 150 Hz (“Thumb” Daily Diary (DD) units, hereafter type 1 tag). One tag was placed on the upper back, the other on the lower back, both in the dorsal mid-line. Units measured 22 by 15 by 9 mm and the distance between them was *ca*. 4 cm.

To ensure that only steady sustained level flight was included in the analysis, we selected sections of consistent flapping flight lasting for at least 2 s (corresponding to *ca*. 10 wingbeat cycles), with no gliding or wingbeat interruptions. The stability of the flight was controlled by selecting sections where VeDBA values smoothed over 1 s were between 0.75 and 3 *g* and varied by less than 1.0 *g*, with no apparent trend (increasing or decreasing) over time. We also discarded the first second of any flight.

We first assessed whether the VeDBA values differed with tag position. VeDBA was calculated using a 2 s smoothing window to derive the “static” component (Shepard et al., 2008) and then subtracting static values from the raw acceleration data in each axis, before summing the differences vectorially (Qasem et al., 2012). We then assessed whether the peak amplitude per wingbeat differed according to tag location, with the peak amplitude calculated as the difference between the maximum and the minimum value of heave acceleration. For this, peaks were detected in the heave axis (Bishop et al., 2015) to synchronise every wingbeat to a defined start point. Finally, to understand which parts of the wingbeat signal were affected by the difference in tag position, we analysed the acceleration signals across average wingbeats in the three acceleration axes. Each acceleration datapoint was attributed to a percentage progression across the wingbeat cycle. Then, for every whole percentage value, the heave, surge and sway accelerations were averaged across 10 wingbeats from the same logger. The average values for the heave, surge and sway accelerations of the upper back-mounted tag were expressed against the values of the lower back-mounted tag in a linear model, the slope of which was used to determine the difference in signal amplitude between the two tags for each acceleration axis.

To examine putative changes in heave signal amplitude (see above) and VeDBA associated with tag placement, we compared them between upper- and lower-back tags using a paired Student’s t-test for VeDBA and a Wilcoxon signed-rank test for amplitude (due to non-homogeneous variances between the two groups (Levene’s Test: F-value = 4.159, p = 0.049)). Wingbeat frequency also contributes to the variation of VeDBA (Van Walsum et al., 2020). Wingbeat frequency was also compared between the two tags using a paired Student’s t-test. The statistical analysis was performed in RStudio, using R version 4.0.3 (R Core Team, 2020).

### Acceleration error in field studies

As a *post hoc* example of how different deployment protocols may affect accelerometer-based results, we compared the amplitude of the heave acceleration signal and VeDBA during the flight of black-legged kittiwakes for two different setups. Twelve kittiwakes were captured and tagged during their breeding season on Middleton Island, Alaska (59.43 N, 146.33 W) and equipped with an accelerometer (type 1 DD) placed under their tail, sealed inside heat shrink tubing for waterproofing: This method is popular as it prevents the bird from trying to preen off the package. We equipped 4 other birds with the same tags placed on their back and wrapped in two zip-lock bags to protect them from splash damage, while allowing pressure sensors to function: This other method is particularly favoured in studies aiming to measure altitude, as it does not require a full waterproofing, which alters pressure recordings. Tail-mounted tags were also tied to a GPS, while the back-mounted units were in an independent package so that the back-mounted logger package was 1 g heavier (total masses; tail = 21 g, back = 22 g). Two 1-min sections of level flapping flight were identified for each tag and deployment. The selection was made based on the altitude data from the loggers’ pressure sensors (< 5 m difference between the highest and lowest altitude measurements), after verifying that there was no interruption in the wingbeat pattern found in heave, ascertaining that the bird flapped regularly for the whole period.

In a similar manner, we examined red-tailed tropicbird data from two different nesting seasons using tags placed in a standard position on their lower back while using different tags. For this, red-tailed tropicbirds at Round Island (19.85 S, 57.79 E) were captured on their nests and equipped with two different units by the same person using 4 strips of Tesa tape placed under the feathers and around the tags (Wilson & Wilson, 1989). Nineteen birds were tagged between February and March 2018 (using type 2 DDs, Figure 1) while 36 birds were tagged during the second season (September and October 2018, type 2 DDs, Figure 1). Importantly, during the second season though, the tags were attached using only 3 strips of tape. At the time, this was considered adequate and helped reduce the weight of the unit. Both units were set to the same sampling frequency (40 Hz). They were however built with different accelerometers (type 1: LSM9DS1, type 2: LSM303DLHC, STMicroelectronics, Geneva, Switzerland), with a substantial difference in sensitivity (type 1: 0.061 *mg*, type 2: 1.0 *mg* sensitivity at +/− 2 *g* range). In addition, the accelerometer is placed at the front of the type 1 unit, and at the back of the type 2 unit, leading to an estimated distance of up to 1 cm between them once placed on the bird’s back. The type 1 tags used in the second season were slightly lighter (masses; type 1 unit = 25.0 g, type 2 unit = 27.7 g). As with the kittiwakes, level flapping flight was selected to discard the effect of gliding, thermal soaring or climbing on acceleration metrics (Williams et al., 2015). We considered level flapping flight to be any section where VeDBA > 0.3 *g* and where the rate of change of altitude (measured by the pressure sensor of the Daily Diary at 4 Hz) was between −0.5 and 0.5 ms^−1^.

**Figure 1:**
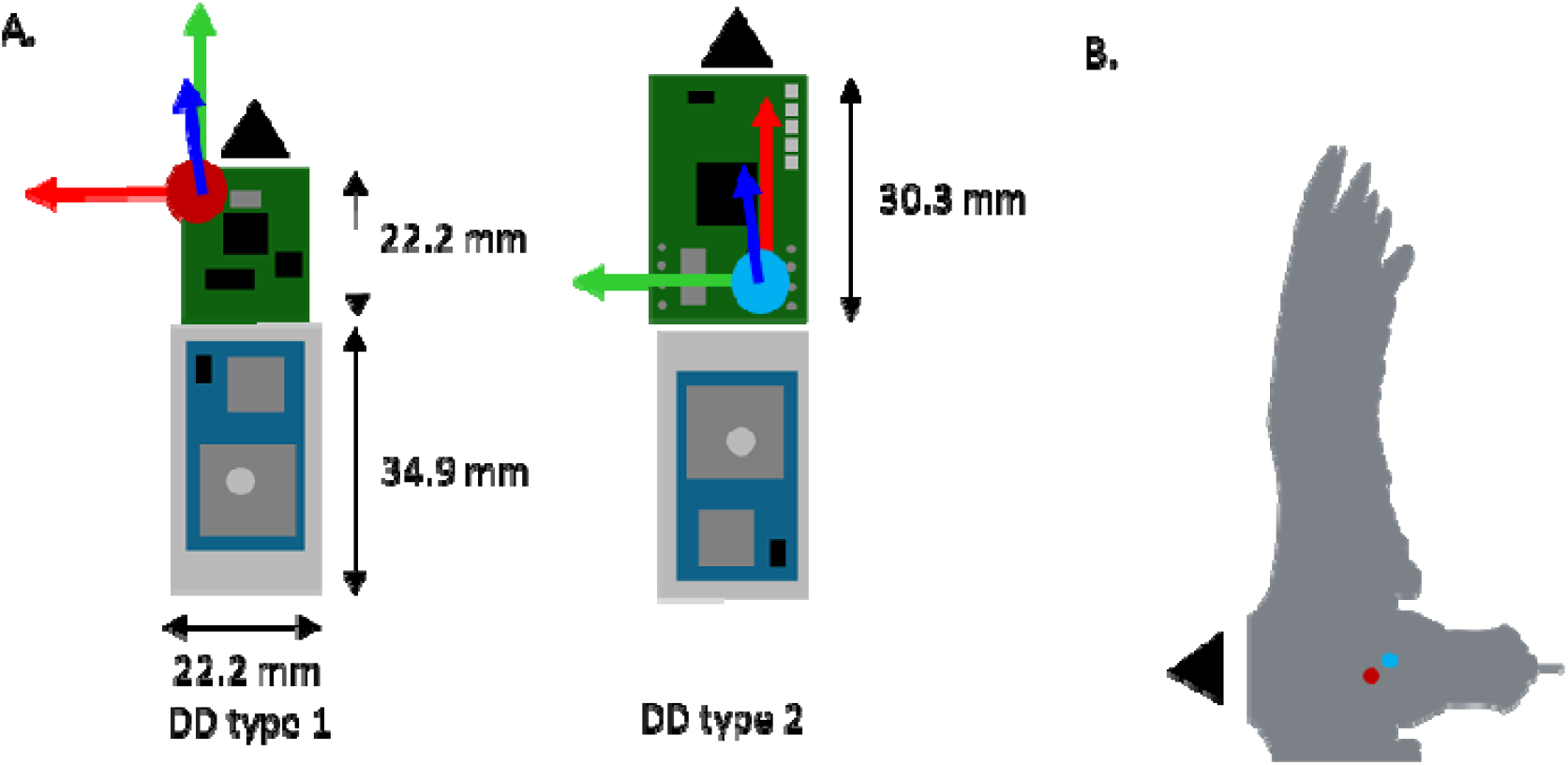
(A) Location of the accelerometer [interception point of the 3 arrows depicting tri-axial acceleration] on the circuit boards of two different DD tags [the battery is in grey, the GPS in blue and the DD in green] and (B) location of the accelerometers within the tags on the back of a red-tailed tropicbird for the type 1- (red dot) and type 2- (blue dot) tags.

VeDBA, wingbeat frequency and the amplitude of heave in level flapping flight were derived from accelerometer data for both tropicbirds and kittiwakes following the same process as pigeons. Data were not paired, since birds carried one tag at a time, so non-paired Student’s t-tests and Wilcoxon tests were used to compare the three parameters between loggers.

Since both the tropicbird and kittiwake data were collected from uncalibrated accelerometers (see above), a situation that we believe represents most of the accelerometer deployments made by the community to date, we attempted to assess the potential for accelerometer error *post hoc*. We did this by measuring the variability in the vectorial sum at times when the tags were motionless (though not on the study animals) and in different tag orientations, finding up to 5 different orientations per logger (for example when units were placed inside bags and the bag placed on the floor/ground). The mean vectorial sum of the three axes of acceleration was calculated for each orientation, and compared between loggers and between tag versions using two ANOVAs.

## Results

### Measurement of acceleration accuracy of tri-axial sensors

Static calibrations of the 15 separate accelerometers within the 5 tags showed that axis offsets needed corrections up to between −0.043 and 0.025 *g* and had multiplicative factors ranging between 0.97 and 1.023. Mean multipliers (across all three axes) for any one tag ranged between 0.9933 and 1.0147.

In the walking speed trials with people, the minimum and maximum differences in VeDBA between calibrated and uncalibrated tags for any one participant ranged between 0.37% and 5.04%. Mean VeDBAs per participant across speeds showed that the difference between calibrated and uncalibrated tags could amount to 2.5% of the calibrated reading. Inspection of the measures undertaken to calibrate each tag (see above) showed that the percentage difference between the uncalibrated and calibrated was primarily due to the acceleration multiplicator (see above) (Figure 2).

**Figure 2.**
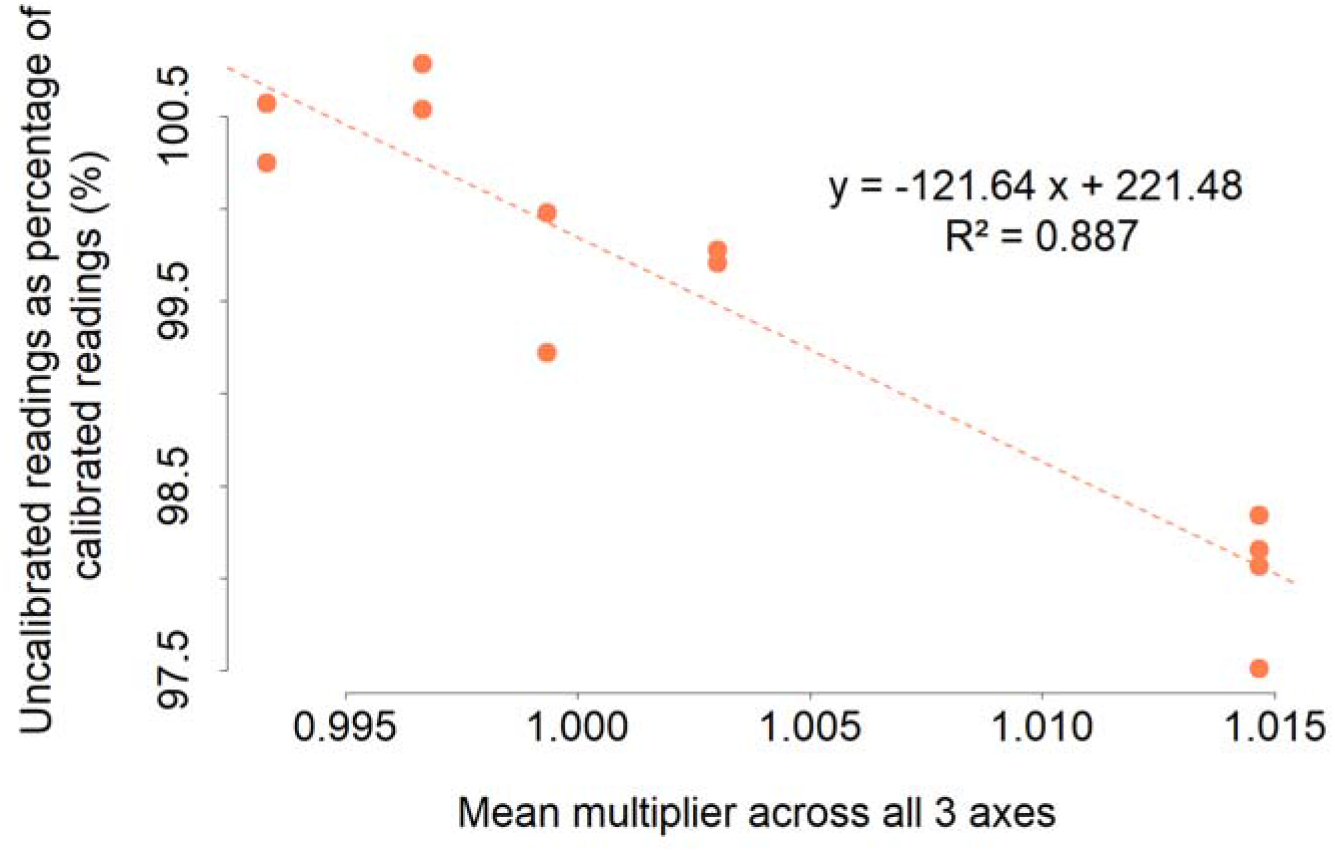
Percentage difference between VeDBA values derived during controlled speed trials with walking humans using uncalibrated against calibrated (corrected) values. The mean multiplier is one applied across all three axes and does not represent the range of values between axes, which can be considerably higher (see text).

### Effect of tag position on raw acceleration in pigeons

In our controlled study with pigeons, plots of surge *versus* heave acceleration showed how wingbeats under identical conditions returned markedly different profiles of acceleration depending on the tag position (Figure 3. A). We also found corresponding differences in values of the heave and surge according to tag position (Figure 3. B, C and D): the upper tag recorded a lower magnitude of surge (LM: Estimate = 0.76, p < 0.001, R^2^ = 0.41, with a slope < 1, Figure 3. C), but a higher magnitude of heave than the lower tag (LM: Estimate = 1.2, p < 0.001, R^2^ = 0.97) (Figure 3. D). The sway model however, showed a weak fit (LM: Estimate = 0.18, p < 0.001, R^2^ = 0.18) and the slope of their relationship was < 1 (Figure 3. B).

**Figure 3:**
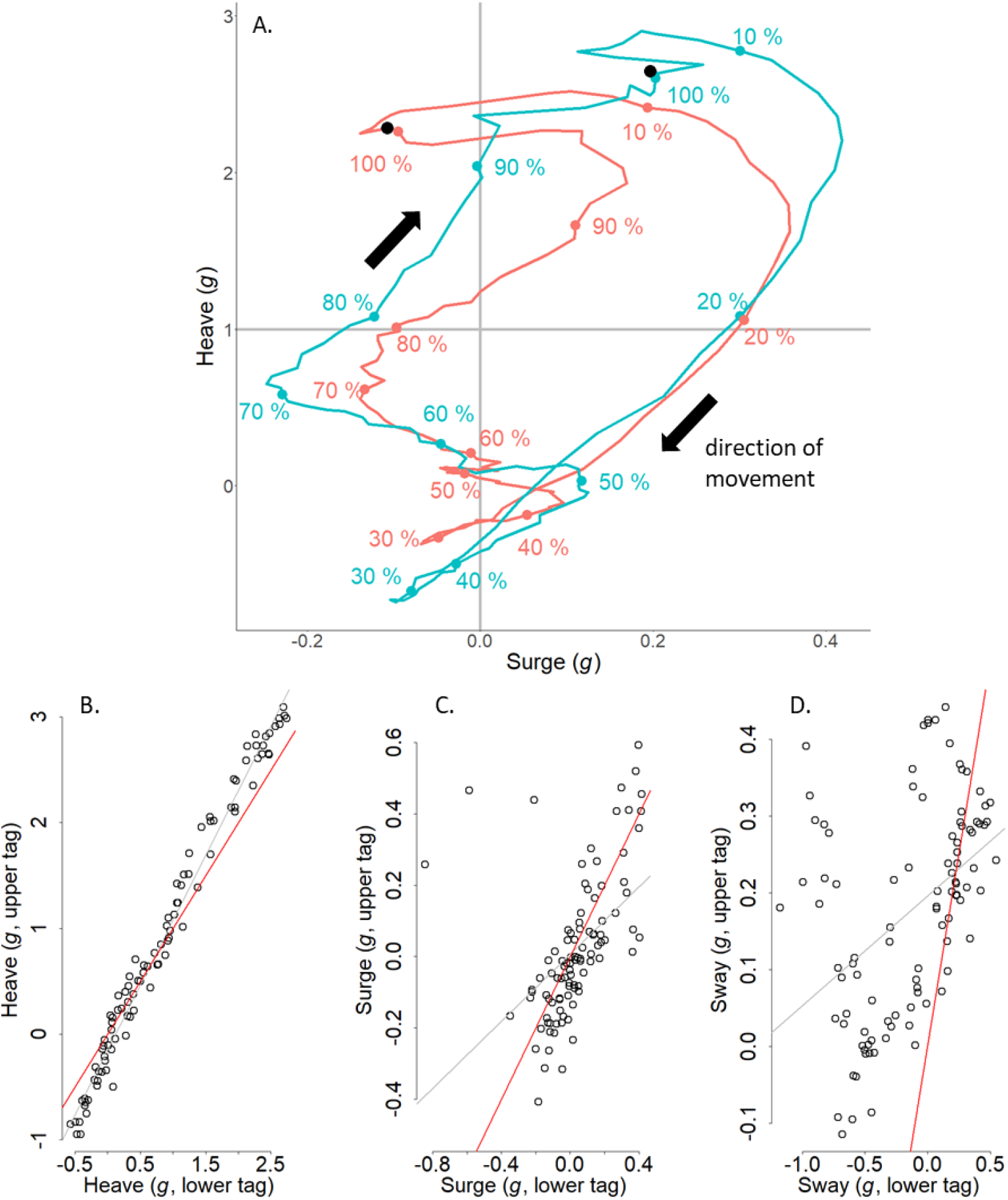
(A) Plot of mean heave versus surge acceleration through time for a pigeon during an average wingbeat cycle derived from a lower back (red) and an upper back-mounted tag (blue), both recording at 150 Hz. Each point corresponds to a mean value of acceleration calculated across all flights for a given percentage through the wingbeat, starting from the peak of acceleration of the downstroke (black point). The value of each point was smoothed over a window of 10 points (10%) to reduce noise. Regressions of the upper against lower tag acceleration for defined points throughout the wingbeat cycle show; (B) heave, (C) surge and (D) sway accelerations (note the changing axis scales). The regression between the two tags is represented in grey, and the y = x line is shown in red.

### Effect of tag position on acceleration metrics

Differences in raw acceleration values also resulted in some variation in acceleration-derived metrics in both the controlled studies on pigeons and in the *post hoc* studies on wild birds: Upper back-mounted tags recorded a slightly higher VeDBA than lower back-mounted tags in pigeons (paired Student’s test: difference = −0.167, t = −2.184, p = 0.043), which was largely due to higher heave values (Wilcoxon signed-rank test: difference = 0.82 *g*, W = 94, p = 0.007) (Figure 4. A, D).

**Figure 4:**
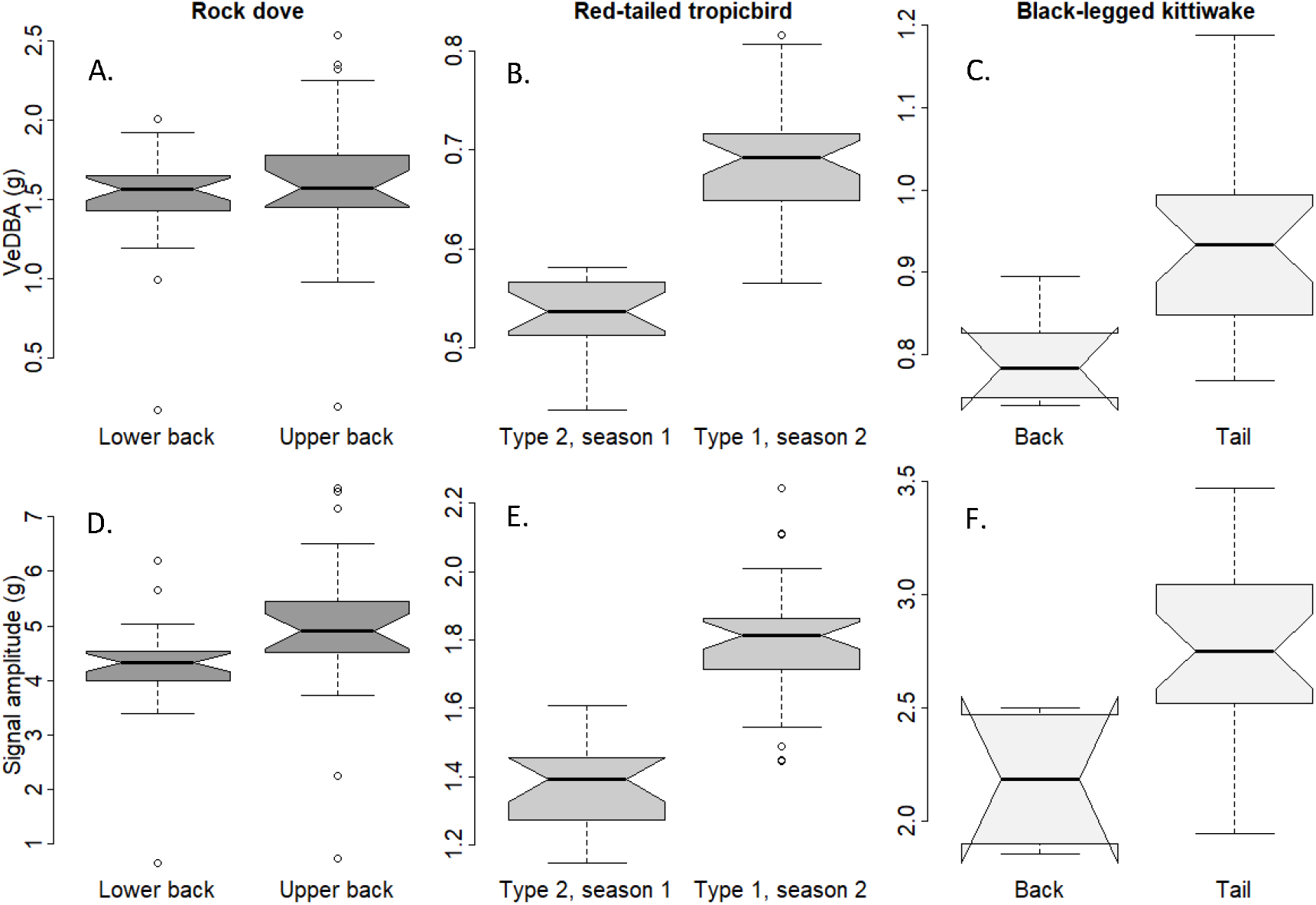
Comparison of VeDBA (A, B, C) and heave signal amplitude (D, E, F) between tags in pigeons (A, D), red-tailed tropicbirds (B, E) and black-legged kittiwakes (C, F). Bold horizontal lines indicate the median vectorial sum for each tag, extremes of the box the upper and lower quartiles, and whiskers the extreme values (excluding outliers, represented by open circles). Notches represent 1.58 IQR/√n (n being the number of observations) on either side of the median and suggest a significant difference when they do not overlap.

In red-tailed tropicbirds, the type 1 tags, used during the second deployment, recorded both a higher VeDBA (by 25%) (Wilcoxon test: difference = 0.14 *g*, W = 19, p < 0.001) and heave amplitude (by 29%) (Student’s t-test: difference = 0.40 *g*, t = −11.78, df = 47.718, p < 0.001) than the type 2 tags (Figure 4. B, E). In kittiwakes, the tail tags recorded both a higher VeDBA (by 18%) (Wilcoxon test: difference = 0.14 *g*, W = 14, p = 0.001), and a higher heave amplitude (by 27%) (Student’s t-test: difference = −0.60 *g*, t = −4.4304, df = 9.0178, p-value = 0.002) than the back-mounted tags (Figure 4. C, F).

There were no differences in estimated wingbeat frequency according to where tags were mounted in either pigeons (paired Student’s t-test: t = 1.954, p = 0.067) or kittiwakes (Wilcoxon test: W = 100, p = 0.227). In tropicbirds, there was a seasonal difference in wingbeat frequency, with type 2 tags recording a higher wingbeat frequency (by 3%) than the type 1 DDs (Student’s t-test: difference = − 0.14 Hz, t = 3.72, df = 35.19, p < 0.001).

We found a positive relationship between wingbeat frequency and heave amplitude during tropicbird level flapping flight (LMM, Season 1: estimate = 0.249, intercept = 0.254, std. error = 0.021, t = 13.339, p < 0.001; Season 2: estimate = 0.746, intercept = −1.084, std. error = 0.024, t = 19.710, p < 0.001; 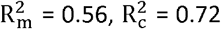). The slope was however steeper during season 2 (Figure 5), in line with the higher amplitude of heave recordings (see Figure 4).

**Figure 5:**
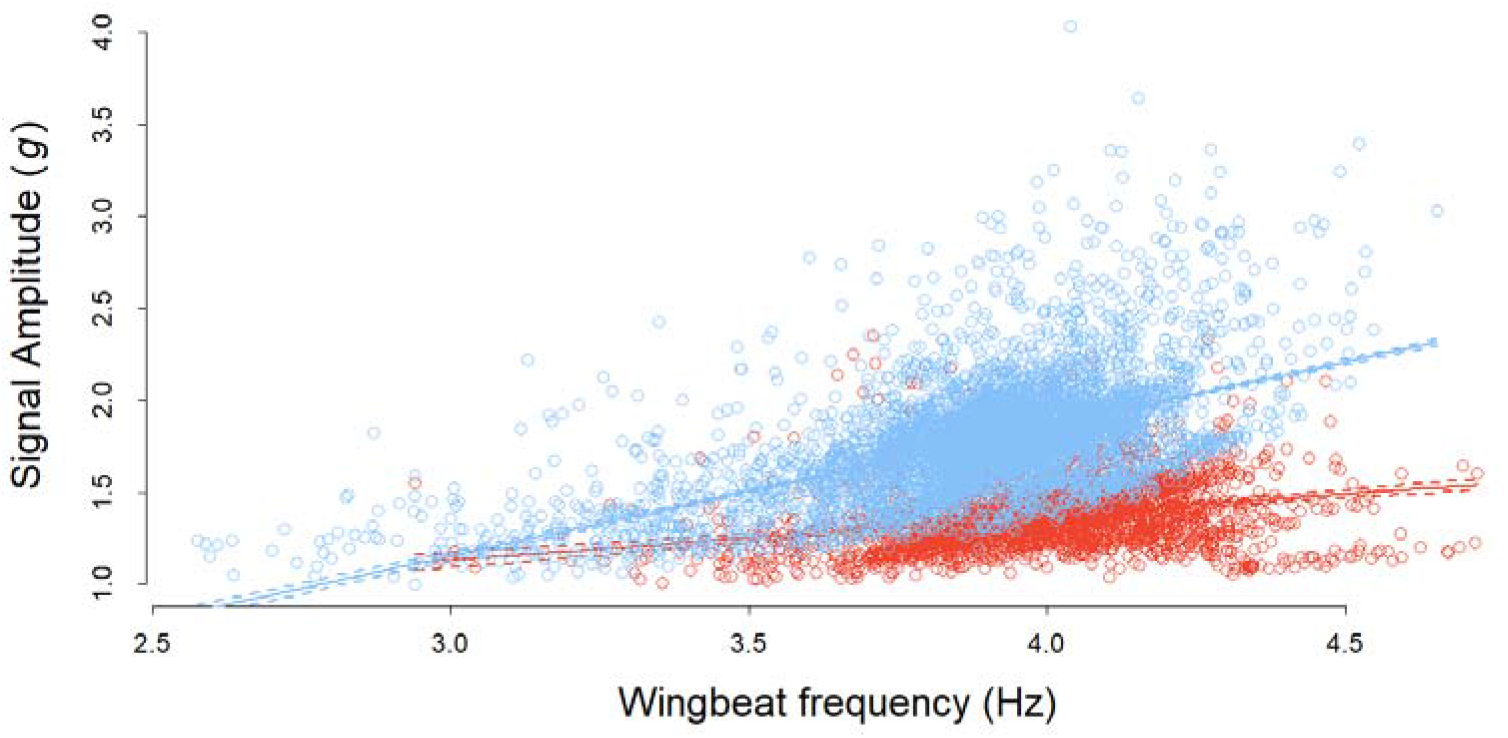
Relationship between the wingbeat frequency and heave amplitude of red-tailed tropicbirds during two field seasons. Birds were equipped with type 2 tags in season 1 (red) and type 1 tags in season 2 (blue), using one less strip of tape, which could reduce tag stability. Full lines represent the linear relationship between wingbeat frequency and amplitude and dashed lines its confidence interval.

### *Post-hoc* quantification of accelerometer inaccuracy

The comparison of stationary data recorded by the two tag types deployed on tropicbirds indicated that the vectorial sum was lower in the type 2 tag (Wilcoxon test: W = 98, p = 0.005, difference = 0.03 *g*) (Figure 6). Standard deviations of the vectorial sum (Type 1: 0.03; Type 2: 0.05) however, indicate that errors are more variable within type 2. We could not determine multipliers for the three acceleration channels to calibrate the data based on this approach, as the heave and surge channels did not cover the whole spectrum of their possible distribution (−1 to 1 *g*) while the tag was motionless.

**Figure 6:**
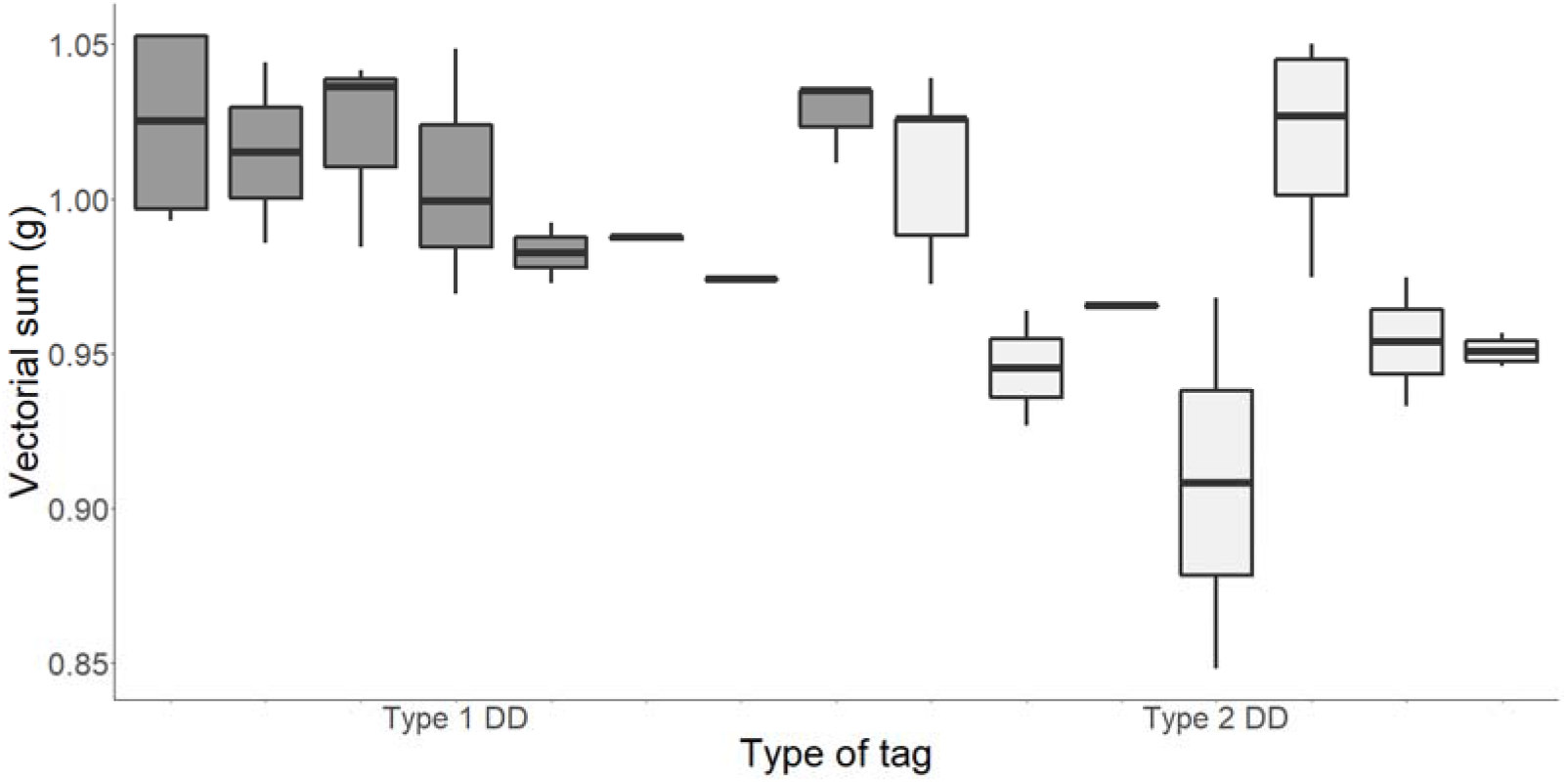
Comparison of the vectorial sum of the raw acceleration recorded by various immobile type 1 (dark boxes) and type 2 tags (light boxes). Each point corresponds to a different unknown orientation. Thick black lines indicate the median vectorial sum for each tag, extremes of the box the upper and lower quartiles, and whiskers the extreme values (excluding outliers).

## Discussion

This work highlights that variation in acceleration measured by tags on flying birds (and presumably other animals engaged in any activity) can be due to; (i) differences in sensitivity (Table SI1) and calibration between sensors, (ii) variation due to the placement of the tag (or the sensor within the tag) and (iii) variation due to the animal itself. Of these, it is normal to attribute all variation to the activity of the animal itself but the validity of doing this is critically dependent on the other two. Studies that do not consider points (i) and (ii) may, therefore, be misrepresenting animal activity both in terms of intensity and extent. We propose here a calibration method to prevent sensor-induced errors, and provide some recommendations about tag attachment method to avoid interpreting tag position effect as biological.

The variation in acceleration is used to examine animal behaviour within a multitude of research thrusts, some of which use acceleration data in slightly different ways. These range from the precise definition of heave, surge or sway values or derivatives (such as pitch and roll and DBA) used in algorithms to identify behaviours (e.g. Fehlmann et al., 2017; Nathan et al., 2012) through the use of acceleration-derived metrics to define energy expenditure (in e.g. doubly labelled water versus DBA regressions (Pagano & Williams, 2019)), to measure travelling speed (Bidder et al., 2012; Gunner et al., 2021) and studies looking at animal effort over time and space (Duriez et al., 2018; Halsey et al., 2011). Errors due to sensor inaccuracy and differences in placement are most severe when axes are considered individually (e.g. deriving pitch from the surge axis). However, they are also relevant when all three orthogonal axes are considered, as inaccuracies in one axis can either be mitigated or compounded by inaccuracies in another (see Figure 2). Within vectorial (or absolute) sums of acceleration metrics, the overall error will depend on the relative errors of the different axes and the extent to which they vary during the activity in question. For example, in flapping birds or bats, almost all variation in acceleration measures occur in the heave and surge axes (e.g. Wilson et al., 2008, and see Figures 3 A-E) so errors in the sway are less important. Cognisance of the axis-specific errors will help mitigate those errors that could be interpreted as a biological effect.

### Calibrations

The issue of inaccurate sensors can be at least partially mitigated by the 6-O method suggested in this work, although we note that this only effectively calibrates between −1 and 1 *g*, while the gravitational component experienced by some animals e.g. during turning (Wilson et al., 2013), will increase beyond these limits. Although, ideally, the tags should be calibrated with each of the accelerometer axes held perfectly vertically (something that is challenging to do once a circuit board is potted in a housing), in practice, this is not critical, and holding the axes as close to vertical as possible should suffice. This is because the response of an accelerometer to the static acceleration of the earth’s gravity follows a sine wave so that an accelerometer that is placed 10° off the vertical (i.e. at 80°), reads a value that is 98.5% of the full-scale value that would be given if the accelerometer axis were held perfectly vertical (so that if there is an error in this axis, 98.5% of it will be covered by this orientation). If it is impossible to reliably estimate the angle of the logger because of the housing, for instance, gently rotating the logger around in every direction would be needed to cover all 6 orientations. Using this calibration will therefore allow researchers to ascribe the most substantive variation in acceleration signal to specific axes.

Our suggestion of dealing with errors *post hoc* by looking at the vectorial sum of the acceleration when tags were stationary could not be used to correct the various axes in our study because all 6 orientations required for the calibrations were not known. However, this process does at least serve to indicate some of the extent of deviation of the sensors from the expected range (see above – Figure 4). In this regard, we note that we have presented results in this work from only one tag manufacturer (type 1 and type 2 tags use two different chips; the type 1 is far superior having a sensitivity of 0.061 mg (in a range of +/− 2 *g*), while the type 2 only has a sensitivity of 1 m*g* for this range), but we have measured, in passing, more substantive variation by other manufacturers (see SI 1).

### Why does accelerometer position affect acceleration?

The position of an accelerometer on an animal should affect the acceleration perceived by the sensor during movement according to its location, and indeed that is the basis behind many biomechanical studies (e.g. Giansanti et al., 2003; Hyde et al., 2008). However, there is poor appreciation in the behavioural ecology community that this premise is also valid for trunk-mounted tags. This may seem irrelevant for birds where the thorax can be considered a single immobile unit, in contrast to bead-string models that may indicate what is expected in species with a flexible back (Underhill & Doyle, 2006). Our work has shown, however, that the location of trunk-mounted accelerometers on birds does play a role in modulating acceleration values (Figure 3) and this is presumably because the bird body pitches during the wingbeat cycle (although part of the differences that we observed may also be due to the movement of the scapulae and perhaps the neck during flapping). Depending on the degree of pitch, the centre of pitch rotation and the position of the accelerometer, this will change the extent of movement (d), which can be defined by the length of a section of a circumference around the centre point of rotation according to D = 2πr(360/P), where r is the radius or distance between the centre of pitch rotation and the sensor, and P is the maximum pitch angle (in degrees). The duration of the wingbeat cycle will define the vertical speed of the tag at its location, with the recorded acceleration being the change in speed over time. The formula shows how the effect of changed acceleration will be manifest more with increasing distance of a tag from the centre point of rotation and so will have the greatest potential to vary in larger birds, all other things being equal. This may also account for the changed acceleration metrics in tail- versus body-mounted tags (Figure 4 C, F) in our kittiwake study although part of that is presumably due to the relative instability of the tail. In fact, to our knowledge, there is little information on the extent of bird body change in pitch during flight (but see Su et al., 2012; Tobalske & Dial, 1996) although controlled experiments with multiple calibrated accelerometers could change that. In the meantime, we suggest that users attempt to place accelerometers in identical positions on their study animals for comparative purposes, which should also involve knowing the position of the sensors within the tags rather than just considering the tags themselves (Figures 1 and 3).

Fortunately, there is no *a priori* reason why tags placed differently on a bird thorax or inaccurate accelerometers should affect determination of wingbeat frequency since points of inflection will still be represented correctly with respect to time within the wingbeat cycle (Figure 3 A, B). Indeed, this is what we observed in our controlled pigeon flight trials and in the kittiwakes (despite a small difference in tag mass, see Whelan et al., 2021). In contrast, the tropicbird work indicates that there was indeed a change in wingbeat frequency across the two seasons, and this seems to be related to changes in environmental conditions (Garde *et al., in prep*).

### *Post-hoc* studies and differences between tags

The bigger question is the extent to which observed differences between conditions in uncalibrated accelerometers can be attributed to the animals rather than to tag position, attachment techniques or sensor variability. In our tropicbird example, the differences in VeDBA and signal amplitude were not consistent with the differences found in pigeons (higher values in the upper tag), suggesting that they were not related to tag position. Importantly, the difference in amplitude was appreciably larger between the type 1 and type 2 tags on tropicbirds, than between the upper and lower tags used in pigeons, even though the tropicbird tags were placed in a way that minimised the distance between their respective accelerometers. The variability in the vectorial sum of the acceleration between orientations of the same tags (Fig. 4) only amounted to an average difference of 3%, which is close to the difference found between tags used for the <6-O calibration (2%). In contrast, in flapping flight, the difference in VeDBA between tags and seasons reached 25%. This order of magnitude difference, coupled with the observation that the difference between vectorial sum values in 6-O calibrated tags and uncalibrated tags (in general) amounted to a mean maximum of 2.5% (similar to VeDBA differences across human walking trials of 2.5%), would appear to indicate that the differences observed in the tropicbird studies were due to seasonal changes in the birds’ interactions with the environment. This is backed up by the changes in wingbeat frequency between the two seasons, which would not be affected by either tag position or sensor inaccuracies. However, Wilson et al. (2021) note how accelerometers on loosely fitted collar tags on mammals provide a signal that effectively depends on collar tightness: Under normal conditions, when the tag is tightly associated with the body, the unit replicates the body movement and accelerations faithfully. However, when the attachment is loose, the tag is projected forward and upward during the initial phase of a stride cycle because the tag (and/or collar) abuts the body. This is followed by a short-term dissociation when the tag is not in proper contact with the body, followed again by substantive acceleration as the body catches up with the tag in the proximate interaction. Importantly, this acceleration is higher than that of the body because the animal body is surging forward and upward again while the tag is falling back so that the recorded acceleration spike mirrors the difference between these two processes. Although the attachment of devices to birds using tape (Wilson et al., 1997) provides a much more intimate association between the tag and the bird body, we believe that if this method is not standardized (and it was not in our study), it can lead to major variation in acceleration values, particularly in animals with highly dynamic movement. In birds, this issue may be exacerbated by tag movement due to air flow over the body which can cause the device to vibrate more or less depending on attachment (cf. Wilson et al., 2020). It is also germane to consider that tag attachment stability may change over time in longer-term deployments. These issues have long been recognised in the wearable sensors industry for humans (Jayasinghe et al., 2019). Consequently, we cannot, in good faith, compare VeDBA or wingbeat amplitudes of tropicbirds between seasons although the wingbeat frequency will be unaffected.

### Conclusions: The importance of calibrating loggers and standardising protocols

Accelerometer inaccuracies can result in errors in the raw acceleration of up to 5% per axis and, depending on the extent and direction of the errors across all three orthogonal axes, this can affect DBA metrics accordingly. Tag placement can also result in errors in DBA metrics of up to 9.7% in flapping flight for our units, although we note that the scale of the errors varies between device types. Finally, non-standardized tag attachment procedures can result in highly variable dynamic acceleration values. Taken together, these represent a potentially important source of error in both raw acceleration values, which are commonly used to calculate body pitch and roll and/or as parameters to define particular behaviours, and derived metrics such as DBA. Attachment procedures should be adapted to the species tagged, as the effect of different tag placements may vary from one species to the other (e.g. Kölzsch et al., 2016; Vandenabeele et al., 2014), and to the study, as different metrics may be measured more reliably using one particular method (Kölzsch et al., 2016), making the use of a standardised procedure difficult. Animal disturbance and study purposes should be considered before adjusting tag placement for the compatibility of datasets, and therefore, researchers should be aware of the attachment methods used to compare acceleration metrics between studies reliably (Sequeira et al., 2021). Importantly, we highlight that sensor inaccuracy can be mitigated by performing a rapid calibration. There is therefore a need for researchers to undertake such calibrations prior to each deployment and include this in their archived data as well as to standardize their tag attachment procedure as much as possible. The last decade has been hailed as a golden age in bio-logging, due to the availability of powerful sensors in animal-attached technologies. The data repositories that archive these data represent extremely valuable resources for the community (e.g. Davidson et al., 2020), but there is an urgent need for calibrations that allow data to be standardized in order for their full potential to be realized now and in the years to come.

## Data, code and materials

Data and code used for the analyses of this manuscript are available from the Dryad Digital Repository.

## Authors’ contributions

Red-tailed tropicbird data was collected by AF, NC and VT. Human data was collected by KARR, RPW and RSM. Pigeon data was collected by HR and ELCS under the supervision of MW. Black-legged kittiwake data was collected by BG, FT, SW and KHE. MDH designed the tags used in this study and provided technical information about accelerometers.

Data analysis was carried out by RPW and BG. Calibrations were designed by RPW and MDH. The manuscript was written by BG, RPW and ELCS. All authors contributed to the revision of the manuscript, gave final approval for publication and agree to be held accountable for the work performed therein.

## Acknowledgements

This work was funded by the European Research Council starter grant 715874 to ELCS, under the European Union’s Horizon 2020 research and innovation program. Special thanks to Jenna Schlener and Lottie Rees-Roderick for their contribution on the kittiwake and the wind tunnel study, respectively, as well as to all the people who participated to the walking experiment.

