## Supplementary material for "Ecological inference using data from accelerometers needs careful protocols": Table SI1

### Supplementary Information

*Table SI 1. Comparison of accelerometer sensitivity between different models. There are a few manufacturers that supply MEMS accelerometers with acceleration ranges of 2, 4, 8, and 16* g*, while the sensitivity, the smallest detectable/measurable change, at different ranges can vary greatly. The table below shows a list of sensors possessing comparable acceleration ranges, the first two of which are used on the devices discussed in this paper, with others from well known manufacturers, in some cases with far less sensitivity than the median. LSM303DLHC is the accelerometer-magnetometer chip built on to type 2 tags, while the LSM9DS1 is the chip built on to type 1 tags. Note that type 1 and type 2 tags have a substantially different sensitivity, far higher in LSM9DS1 accelerometers.*

| **Manufacturer** | **Sensor** | **±2 g** | **±4 g** | **±8 g** | **±16 g** |
| --- | --- | --- | --- | --- | --- |
| **ST** | **LSM303DLHC** | 1 | 2 | 4 | 12 |
| **ST** | **LSM9DS1** | 0.061 | 0.122 | 0.244 | 0.732 |
| **ST** | **MIS2DH** | 0.98 | 1.95 | 3.91 | 11.72 |
| **TDK** | **ICM-20948** | 0.061 | 0.122 | 0.244 | 0.488 |
| **TDK** | **IAM-20381** | 0.061 | 0.122 | 0.244 | 0.488 |
| **TDK** | **IIM-42652** | 0.061 | 0.122 | 0.244 | 0.488 |
| **Analog** | **ADXL346** | 0.015 | 7.81 | 15.63 | 31.25 |
| **Kionix** | **KX132-1211** | 0.061 | 0.122 | 0.244 | 0.488 |
| **Bosch Sensortec** | **BMA180** | 0.244 | 0.488 | 0.977 | 1.953 |
| **Bosch Sensortec** | **BMA456** | 0.061 | 0.122 | 0.244 | 0.488 |
|  | ***Median*** | 0.061 | 0.122 | 0.244 | 0.488 |
|  |  |  |  |  | *mg/LSB* |
